# Mechanics of Live Cell Elimination

**DOI:** 10.1101/2021.08.17.456649

**Authors:** Siavash Monfared, Guruswami Ravichandran, José E. Andrade, Amin Doostmohammadi

## Abstract

Cell layers eliminate unwanted cells through the extrusion process, which underlines healthy versus flawed tissue behaviors. Although several biochemical pathways have been identified, the underlying mechanical basis including the forces involved in cellular extrusion remain largely unexplored. Utilizing a phase-field model of a three-dimensional cell layer, we study the interplay of cell extrusion with cell-cell and cell-substrate interactions, in a monolayer. Independent tuning of cell-cell versus cell-substrate adhesion forces in the model reveals that a higher cell-substrate adhesion leads to a lower number of total extrusion events. We find extrusion events to be linked to both half-integer topological defects in the orientation field of the cells and to five-fold disclinations in cellular arrangements. We also show that increasing the relative cell-cell adhesion forces translates into a higher likelihood for an extrusion event to be associated with a five-fold disclination and a weaker correlation with +1/2 topological defects. We unify our findings by accessing mechanical stress fields: an extrusion event acts as a mechanism to relieve localized stress concentration.

---

The ability of cells to self-organize and to collectively migrate drive numerous physiological processes including tissue morphogenesis [1], epithelial-mesenchymal transition [2], wound healing [3], tumor progression [4] and cancer invasion [5]. Advanced experimental techniques have linked this ability to mechanical interactions between cells [6–8]. Specifically, cells actively coordinate their movements through mechanosensitive adhesion complexes at the cell-substrate interface and cell-cell junctions. In this vein, cell-cell and cell-substrate adhesion seem to be coupled [9], further complicating the interplay of mechanics with biochemistry. Additionally, cell elasticity affects spatiotemporal protein dynamics on cell membrane [10] which consequently affects cellular processes such as cell division [11] and motility [12].

While advances in experimental techniques are followed by more nuanced theoretical and computational developments, a majority of current approaches to simulate multicellular layers are limited to two-dimensional systems, hindering in-depth exploration of intrinsically three-dimensional nature of the distinct forces that govern cell-cell and cell-substrate interactions. Moreover, some of the most fundamental processes in cell biology such as cell extrusion - responsible for tissue integrity - are inherently three-dimensional. Thus, studying the underlying mechanisms necessitates access to both in-plane and out-of-plane features of the cell layers.

Cell extrusion refers to the process of removal of excess cells to prevent accumulation of unnecessary or pathological cells. This process can get initiated through apoptotic signaling [13], oncogenic transformation [14] and overcrowding of cells [15–17] and mediated by actin organization [18]. Most importantly, cell extrusion plays an important role in developmental [19], homeostatic [16] and pathological processes [20], including cancer metastasis. However, the underlying mechanisms that facilitate cell extrusion are still unclear [21].

The similarities between cellular systems and liquid crystals manifested in local nematic alignment and topological defects [22–27] provide a fresh perspective for understanding cellular processes. In cellular monolayers, comet-shapes and trefoils topological defects, corresponding to +1/2 and −1/2 charges, respectively, are prevalent [28, 29]. These are singular points in cellular alignment that mark the breakdown of orientational order [30]. Recent experiments on epithelial monolayers found a strong correlation between extrusion events and the position of a subset of +1/2 defects [22]. Additionally, a numerical study of overlapping active particles in two-dimensions suggest an alternative to topological defects of nematic order, arguing that overlaps between the cells and cellular extrusions are favored near five-fold disclinations [31], representing imperfections in hexagonal, honey-comb structure of cells arrangement in the monolayer [32]. These recently introduced purely mechanical routes to cell extrusion have opened the door to new questions on the nature of forces that are involved in eliminating cells from the monolayer and challenge the purely biological consensus that an extruding cell sends a signal to its neighbor that activates its elimination process [13]. Nevertheless, it is not clear if these different mechanisms are related, and whether, depending on the mechanical features of the cells, the cell layers actively switch between different routes to eliminate the unwanted cells. Since all the existing studies so far have only focused on effective two-dimensional models of the cell layers, fundamental questions about the three-dimensional phenomenon of cell extrusion and its connection to the interplay between cell-generated forces at the interface between cells and the substrate, with multicellular force transmission across the cell layer, remain unanswered.

In this Letter, we explore three-dimensional collective cell migration in cellular monolayers. Based on large scale simulations, we examine (i) the underlying mechanisms responsible for live cell extrusion, including any correlations with ±1/2 topological defects and five-fold disclinations, and (ii) the interplay of cell-cell and cell-substrate adhesion with extrusion events in cellular systems. We find a lower likelihood of an extrusion event occurring as cell-substrate adhesion increases. Furthermore, we link the extrusion events to both half-integer topological defects and five-fold disclinations with the likelihood of an extrusion event associated with each altered by the relative cell-cell interactions. Lastly, by mapping the mechanical stress field across the entire monolayer, we identify localized stress concentration as the unifying factor that governs various mechanical routes to live cell extrusion.

We consider a cellular monolayer consisting of *N* = 400 cells on a substrate with its surface normal 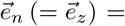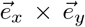 and periodic boundaries in both 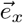 and 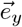, where 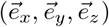 constitute the global orthonormal basis (Fig. 1). Cells are initiated on a two-dimensional simple cubic lattice. The cell-cell and cell-substrate interactions have contributions from both adhesion and repulsion, in addition to self-propulsion forces associated with cell polarity. To this end, each cell *i* is modeled as an active deformable droplet in three-dimensions using a phasefield, 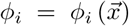. The interior and exterior of cell *i* corresponds to *ϕ_i_* = 1 and *ϕ_i_* = 0, respectively, with a diffusive interface of length *λ* connecting the two regions and the midpoint, *ϕ_i_* = 0.5, delineating the cell boundary. A simple free energy functional, 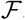, is considered [33]:

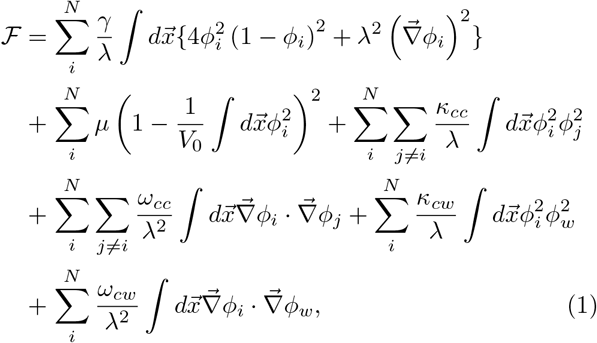

where 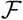 contains a contribution due to the Cahn-Hilliard free energy [34] which stabilizes the cell ifnacte,-followed by a soft constraint for cell volume around 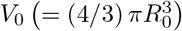, such that cells - each initiated with radius *R*_0_ - are compressible. Additionally, *κ* and *ω* capture repulsion and adhesion between cell-cell (subscript *cc*) and cell-substrate (subscript *cw*), respectively. Moreover *γ* sets the cell stiffness and *μ* captures cell compressibility and *ϕ_w_* denotes a static phase-field representing the substrate. This approach resolves the cellular interfaces and provides access to intercellular forces. The dynamics for field *ϕ_i_* can be defined as:

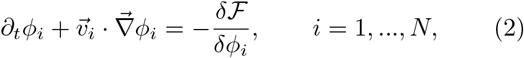

where 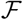 is defined in Eq. (1) and 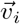 is the total velocity of cell *i*. To resolve the forces generated at the cellular interfaces, we consider the following over-damped dynamics for cells:

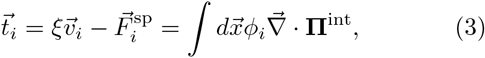

where 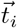 denotes traction, *ξ* is substrate friction and 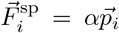 represents self-propulsion forces due to polarity, constantly pushing the system out-of-equilibrium. In this vein, *α* characterizes the strength of polarity force and 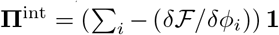. While only passive interactions are considered here, active nematic interactions can be readily incorporated in this framework [9, 33]. To complete the model, the dynamics of cell polarity is introduced based on contact inhibition of locomotion (CIL) by aligning the polarity of the cell to the direction of the total interaction force acting on the cell [35]. The polarization dynamics is given by [36]:

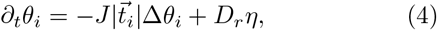

where *θ_i_* ∈ [ −*π, π*] is the angle associated with polarity vector, i.e. 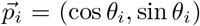 and *η* is the Gaussian white noise with zero mean, unit variance, *D_r_* denotes rotational diffusivity and Δ*θ_i_* represents the angle between 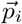 and 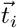. Furthermore, positive constant *J* sets the time scale associated with the alignment of polarity to the total interaction force.

**FIG. 1.**
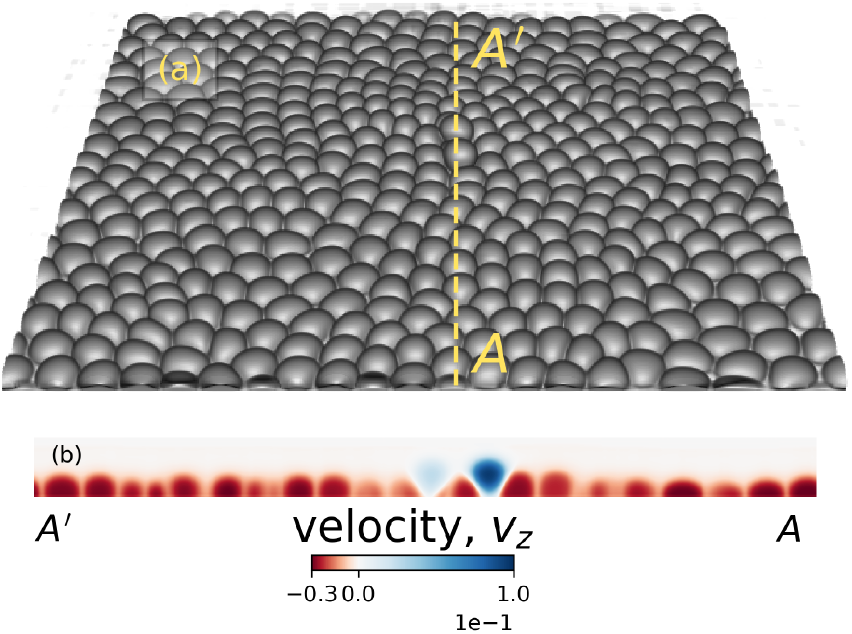
(a): A representative simulation snapshot of a three-dimensional cell monolayer and cell extrusion events. (b): A cross-section (dotted yellow line) of the cell monolayer highlighting extrusion events through out-of-plane velocity 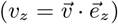.

We perform large scale simulations with a focus on the interplay of cell-cell and cell-substrate adhesion strengths on collective cell migration and its impact on cell expulsion from the monolayer. Following [33], the space and time discretization in our simulations are based on the average radius of MDCK cells, ~ 5*μm*, velocity ~ 20*μm/ h* and average pressure of ~ 100Pa, measured experimentally in MDCK monolayers [22], corresponding to Δ*x* ~ 0.5*μm*, Δ*t* ~ 0.1*s* and Δ*F* ~ 1.5nN for force. In this study, we set the cell-substrate adhesion strength *ω_cw_* ∈ {0.0015, 0.002, 0.0025} and vary the cell-substrate to cell-cell adhesion ratio in the range Ω = *ω_cc_/ω_cw_* ∈ {0.2, 0.4, 0.6}. Based on previous experimental and theoretical studies [9, 33, 36], the other simulation parameters are *κ_cc_* = 0.5, *κ_cw_* = 0.15, *ξ* = 1, *α* = 0.5, *λ* = 3, *μ* = 45, *D_r_* = 0.01 and *J* = 0.005, unless stated otherwise.

In the absence of self-propulsion forces, the initial configuration equilibrates into a hexagonal lattice. Biological processes such as cell division and apoptosis are known to push the system away from a hexagonal configuration [37]. Similarly here, as we introduce self-propulsion forces associated with cell polarity, the system is pushed away from its equilibrium hexagonal configuration, resulting in defects manifested as five-fold and seven-fold disclinations. Fig. 1(a) shows a simulation snapshot with two extrusion events taking place. An extrusion event is detected if the vertical displacement of a cell, relative to other cells in the monolayer, exceeds *R*_0_/2. Fig. 1(b) displays the out-of-plane velocity profile, 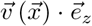 for each cell *i* in the cross-section, clearly marking the extruding cells as they get expelled from the monolayer and lose contact with the substrate.

In order to probe the possible mechanical routes to cell extrusion, we begin with characterizing topological defects in cell orientation field and disclinations in cellular arrangements. To this end, we first map the orientation field of the cells for the entire cell layer from the 2D projected cell shape profile on the *xy*–plane and identify topological defects as the singularities in the orientation field. The results show the continuous emergence of half-integer (±1/2), nematic, topological defects that spontaneously nucleate in pairs and follow chaotic trajectories before annihilation, reminiscent of active turbulence in continuum theories of active nematics [38, 39] (Fig. 2a). It is noteworthy that unlike previous studies of active nematic behavior in 2D cell layers [33, 40], the nematic defects here emerge in the absence of any active dipolar stress or subcellular fields, as the only active driving in these simulations is the polar force that the cells generate. Therefore, although the cells are endowed with polarity in terms of their self-propulsion, the emergent symmetry in terms of their orientational alignment is nematic, which is inline with experimental observations in cell monolayers [22, 25], discrete models of self-propelled rods [41, 42], and recently proposed continuum model of polar active matter [43]. Remarkably, in accordance with experimental observations [22], we find that the extrusion events can be correlated with the position of both +1/2 comet-shaped and 1/2 trefoil-shaped topological defects. To quantify this, Fig. 2(c)-(d) display the probability density of the minimum distance *d*_min_ between an extruding cell at time *t_e_* and ±1/2 topological defects in the interval *t* ∈ [*t_e_* − 900*, t_e_* +100] for four distinct realizations and for varying cell-substrate to cell-cell adhesion ratios Ω. In both cases, the probability density peaks in the vicinity of the eliminated cell (≈ 1.5*R*_0_) and falls off to nearly zero for 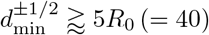

**FIG. 2.**
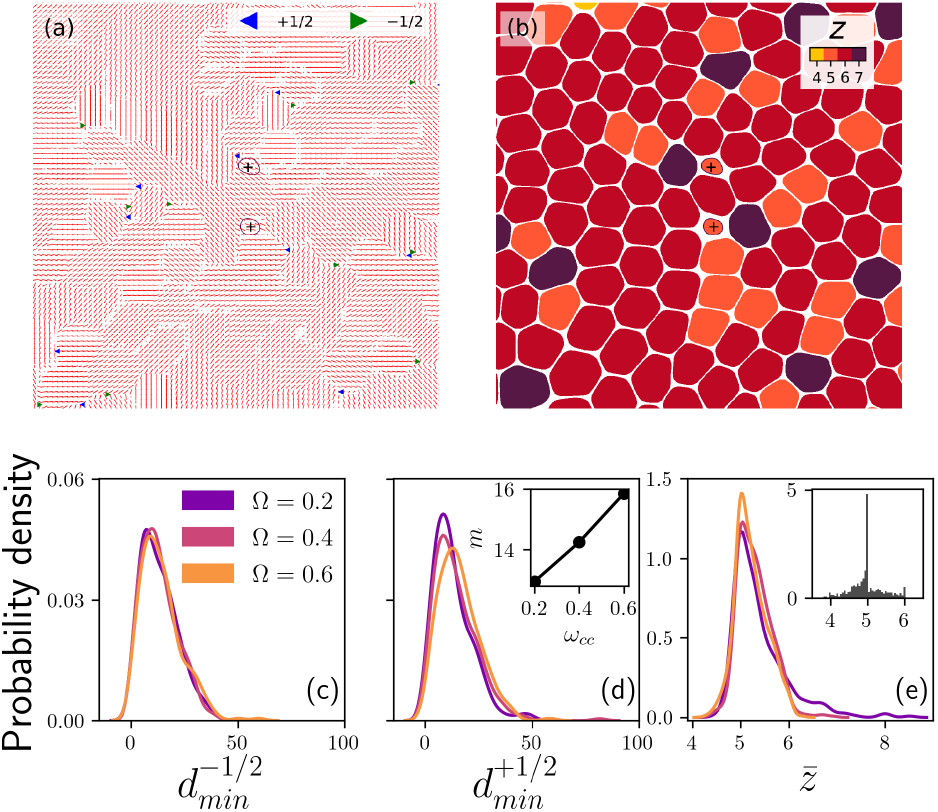
Projected simulation snapshot into xy-plane with (a) ±1/2 topological defects and (b) five-fold and seven-fold disclinations mapped into the monolayer as two extrusion events take place. Probability densities of *d*_min_ for varying cell-substrate to cell-cell adhesion ratios Ω and for four distinct realizations for (c) − 1/2 and (d) +1/2 topological defects (inset: distribution mean *m* vs. *ωcw*). (e) The probabil-ity density of coordination number 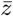 for varying cell-substrate to cell-cell adhesion ratios Ω (inset: 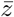 for all extrusion events).

Next, we explore the other possible mechanical route to cell extrusion based on the disclinations in cellular arrangement [31]. To this end, we compute the coordination number of each cell based on their phase-field interactions and identify the five-fold and seven-fold disclinations, with an example shown in Fig. 2(b). To quantify the relation between extrusion events and the disclinations, the probability density of the coordination number of an extruding cell averaged over the same interval, *t* ∈ [*t_e_* − 900*, t_e_* + 100], 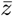, for all the realizations is shown in the inset of Fig. 2(e), clearly exhibiting a sharp peak near 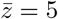.

This correlation between disclinations and extrusion events is also related to the mechanical stress localization at the five-fold disclinations: The occurrence of disclinations in a flat surface produces local stress concentration [32]. Since it is energetically favorable to bend a flat surface, rather than to compress or to stretch it, the local stress concentration can lead to either a five-fold (positive Gaussian curvature) or a seven-fold (negative Gaussian curvature) disclination. In our set-up and given that we consider a rigid substrate, five-fold disclinations are much more likely to provide relief for the high local stress concentration. This can change if the rigidity of substrate is relaxed or extrusion in three-dimensional spheroids are considered. Since we conjecture that both topological defect- and disclination-mediated extrusion mechanisms are closely linked with stress localization, we characterize the in-plane and out-of-plane stresses associated with the simulated monolayer. We compute a coarse-grained stress field [44, 45], 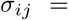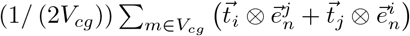 where 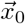 represents the center of the coarse-grained volume,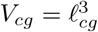 corresponding to coarse-graining length 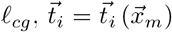 and unit vector 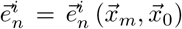 Fig. 3(c) shows the evolution of local isotropic stress, *σ*^iso^ = (1/3) tr***σ***, and an out-of-plane component of the shear stress, *σ_xz_*, aver-aged over an extruding cell *i* and its nearest-neighbors, *nn*. The result shows a clear stress build up, followed by a drop near *t* − *t_e_* = 0, where *t_e_* is detected by our stress-independent criterion. This drop in stress is followed with another build-up for *t* > *t_e_*, due to another extrusion occurring in that vicinity, as highlighted in Fig. 3(b).

**FIG. 3.**
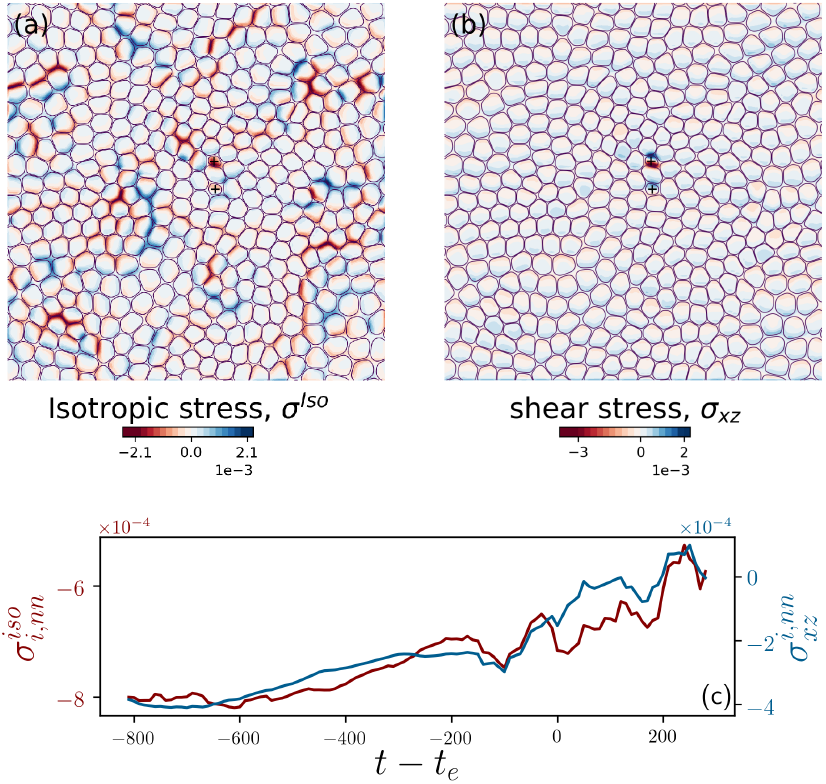
(a): The isotropic stress *σ*^iso^ and (b): out-of-plane shear stress *σ_xz_* fields projected into a plane, with extruding cells marked. (c): Temporal evolution of local averages for *σ*^iso^ and *σ_xz_* for one of the extruded cells. *t_e_* indicates the extrusion time and *R*_0_*/l_cg_* = 4.

Furthermore, we visualize 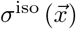 and 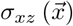 fields at the onset of extrusion events, as shown in Fig. 3(a)-(b). We observe a high, out-of-plane, shear stress concentration as shown in Fig. 3(b) as well as tensile and compressive stress pathways (Fig. 3(a)) reminiscent of force chains in granular systems [46]. While exploring the analogy of the active force chains observed here with those in granular systems is beyond the focus of the current letter, we note that the stress pathways manifest strong correlations with beads of disclination dipoles, i.e., pairs of five-fold and seven-fold disclinations. Interestingly, the association of cell extrusion events with regions of high out-of-plane shear stress has parallels with the phenomenon of *plithotaxis*, where it was shown that cells collectively migrate along the orientation of the minimal in-plane intercellular shear stress [47]. In this context, we conjecture that high shear stress concentration hinders mechanism to re-establish the status-quo.

Next, in order to explore the possible impact of the interplay between cell-substrate and cell-cell interactions, we independently vary the strengths of cell-cell and cell-substrate adhesion and quantify their impact on extrusion events. Figures 4(a)-(b) summarizes these results for four distinct realizations of the considered parameters for cell-substrate adhesion *ω_cw_* and cell-cell adhesion strengths *ω_cc_*. The results show that increasing cell-substrate adhesion leads to less extrusion events (Fig. 4(a)), while the ratio Ω = *ω_cc_/ω_cw_* does not seem to play a significant role on the likelihood of an extrusion event occurring. To explore this further, the temporal evolution of mean and variance of coordination numbers are shown in Figs. 4(c)-(d) for a subset of our simulations. In most but not all cases, increasing cell-substrate adhesion leads to a relatively higher average coordination number temporal evolution with a lower variance while increasing cell-cell adhesion has the opposite effect. This has direct implications on the probability of occurrence of a five-fold disclination and it suggests competing forces that often edge towards cell-substrate adhesion. How-ever, the relative cell-cell adhesion, Ω, seem to alter the likelihood of an extrusion event being associated with a +1/2 defect or five-fold disclination. As shown in Fig. 2(d), the distribution mean for 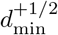, *m*, increases with Ω = *ω_cc_/ω_cw_* (see inset) while the peak of the probability density decreases with Ω. At the same time, as shown in Fig. 2(e), the probability of an extrusion occurring at a five-fold disclination increases with Ω. Together, these results suggest that as the cell-substrate to cell-cell adhesion ratio Ω increases, the likelihood of an extrusion event associated with +1/2 topological defects decreases while the likelihood of such event occurring at a five-fold disclination increases.

**FIG. 4.**
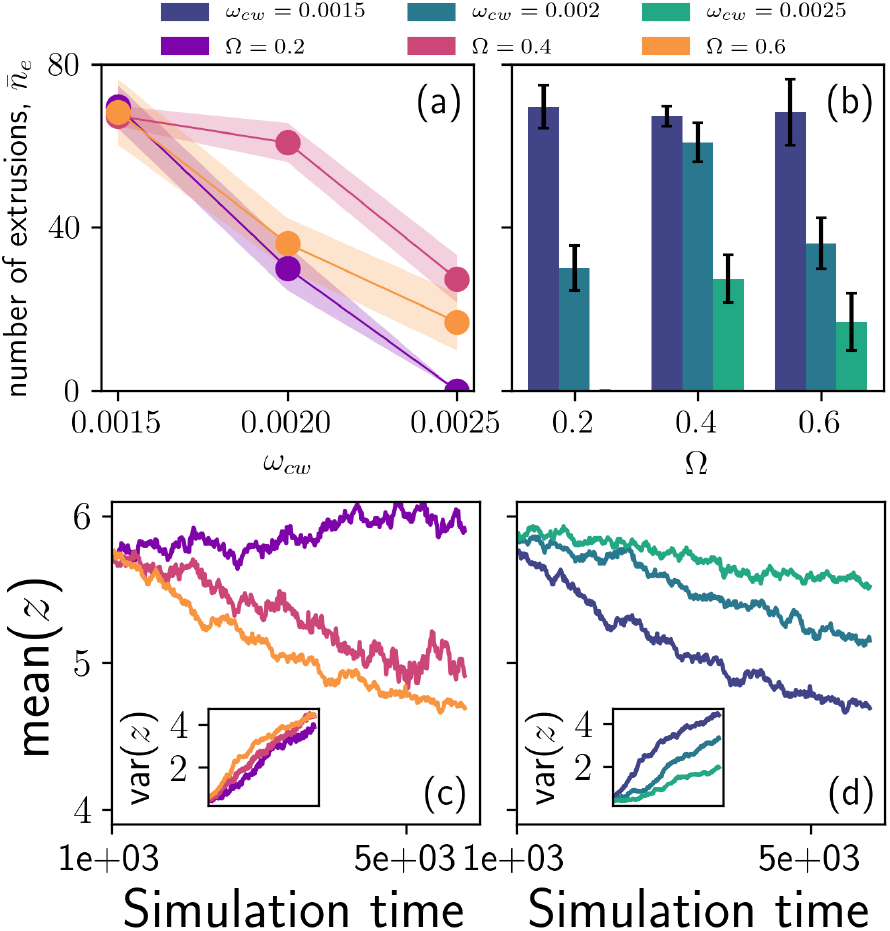
Number of cell extrusions versus (a) cell-substrate adhesion strength *ω_cw_* for fixed values of the cell-substrate to cell-cell adhesion ratio Ω and (b) the adhesion ratio Ω. Temporal evolution of mean and variance (inset) of coordination numbers, *z*, for (c) different Ω and (d) different *ω_cw_*.

Our study presents, to the best of our knowledge, the first three-dimensional model of the collective migration-mediated live cell elimination. Importantly, this framework allows for cell-substrate and cell-cell adhesion forces to be tuned independently. Our findings indeed suggest that varying the relative strength of cell-cell and cell-substrate adhesion can allow cells to switch between distinct mechanical pathways to eliminate unwanted cells through: (i) cell extrusion at ±1/2 topological defects in the cell orientation field, consistent with experimental observations [22]; and (ii) cell extrusion at five-fold disclinations in cell arrangement, where complementing the previous two-dimensional predictions of elevated cell-cell overlaps near disclinations, our results show a direct role of these disclinations in extruding the cells. Focusing on the extruded cells, the results demonstrate that increasing relative cell-cell adhesion increases the probability of an extruded cell being a five-fold disclination while weakening the correlation with +1/2 topological defects. Additionally, the presented framework provides access to the local stress field, including the out-of-plane shear components. Access to this information led us to conjecture that high shear stress concentration frustrates collective cell migration with cell extrusion providing a pathway to re-establish the status-quo, analogous to the *plithotaxis* phenomenon. We expect these results to trigger further experimental studies of the mechanical routes to live cell elimination and probing the impact of tuning cell-cell and cell-substrate interactions, for example by molecular perturbations of E-cadherin adhesion complexes between the cells and/or focal adhesion between cells and substrates, as performed recently in the context of topological defect motion in cell monolayers [9].

Furthermore, we anticipate that this modeling framework opens the door to several interesting and unresolved problems in studying three-dimensional features of cell layers. In particular, the mechanics can be coupled with biochemistry to study a wider range of mechanisms that affect live cell elimination. Additionally, using our framework the substrate rigidity can be relaxed in the future studies to further disentangle the impacts of cell-substrate adhesion from substrate deformation due to cell generated forces. Similarly three-dimensional geometries, such as spheroids or cysts can be examined. Lastly, the links between collective cell migration and granular physics, in terms of force chains and jamming transition, as well as probing the impact of three-dimensionality and out-of-plane deformations on these processes, is an exciting route for future studies.

## ACKNOWLEDGEMENT AND FUNDING

S.M., G.R. and J.A. acknowledge support for this research provided by US ARO funding through the Multidisciplinary University Research Initiative (MURI) Grant No. W911NF-19-1-0245. A.D. acknowledges support from the Novo Nordisk Foundation (grant no. NNF18SA0035142), Villum Fonden (grant no. 29476), funding from the European Union’s Horizon 2020 research and innovation program under the Marie Sklodowska-Curie grant agreement no. 847523 (INTERACTIONS). The authors would like to thank Dr. Lakshmi Balasubramaniam and Prof. Benoît Ladoux (Institut Jacques Monod, Université de Paris), Guanming Zhang and Prof. Julia M. Yeomans (The Rudolf Peierls Centre for Theoretical Physics, University of Oxford), Prof. Jörn Dunkel (Mathematics Department, MIT) and Prof. M. Cristina Marchetti (Department of Physics, University of California Santa Barbara) for helpful discussions.

## References

[1] K. Chiou and E.-M. S. Collins, Why we need mechanics to understand animal regeneration, Developmental Biology 433, 155 (2018).

[2] E. H. Barriga, K. Franze, G. Charras, and R. Mayor, Tissue stiffening coordinates morphogenesis by triggering collective cell migration in vivo, Nature 554, 523 (2018).

[3] A. Brugués, E. Anon, V. Conte, J. H. Veldhuis, M. Gupta, J. Colombelli, J. J. Munõz, G. W. Brodland, B. Ladoux, and X. Trepat, Forces driving epithelial wound healing, Nature Physics 10, 683 (2014).

[4] C. D. Pascalis and S. Etienne-Manneville, Single and collective cell migration: The mechanics of adhesions, Molecular Biology of the Cell 28, 1833 (2017).

[5] P. Friedl, J. Locker, E. Sahai, and J. E. Segall, Classifying collective cancer cell invasion, Nature Cell Biology 14, 777 (2012).

[6] S. A. Maskarinec, C. Franck, D. A. Tirrell, and G. Ravichandran, Quantifying cellular traction forces in three dimensions, Proceedings of the National Academy of Sciences 106, 22108 (2009).

[7] B. Ladoux, Cells guided on their journey, Nature Physics 5, 377 (2009).

[8] B. Ladoux and R.-M. Mège, Mechanobiology of collective cell behaviours, Nature Reviews Molecular Cell Biology 18, 743 (2017).

[9] L. Balasubramaniam, A. Doostmohammadi, T. B. Saw, G. H. N. S. Narayana, R. Mueller, T. Dang, M. Thomas, S. Gupta, S. Sonam, A. S. Yap, Y. Toyama, R.-M. Mège, J. M. Yeomans, and B. Ladoux, Investigating the nature of active forces in tissues reveals how contractile cells can form extensile monolayers, Nature Materials 10.1038/s41563-021-00919-2 (2021).

[10] M. C. Wigbers, T. H. Tan, F. Brauns, J. Liu, S. Z. Swartz, E. Frey, and N. Fakhri, A hierarchy of protein patterns robustly decodes cell shape information, Nature Physics 10.1038/s41567-021-01164-9 (2021).

[11] M. Théry and M. Bornens, Cell shape and cell division, Current Opinion in Cell Biology 18, 648 (2006).

[12] A. Mogilner and K. Keren, The shape of motile cells, Current Biology 19, R762 (2009).

[13] J. Rosenblatt, M. C. Raff, and L. P. Cramer, An epithelial cell destined for apoptosis signals its neighbors to extrude it by an actin- and myosin-dependent mechanism, Current Biology 11, 1847 (2001).

[14] C. Hogan, S. Dupré-Crochet, M. Norman, M. Kajita, C. Zimmermann, A. E. Pelling, E. Piddini, L. A. Baena-López, J.-P. Vincent, Y. Itoh, H. Hosoya, F. Pichaud, and Y. Fujita, Characterization of the interface between normal and transformed epithelial cells, Nature Cell Biology 11, 460 (2009).

[15] E. Marinari, A. Mehonic, S. Curran, J. Gale, T. Duke, and B. Baum, Live-cell delamination counterbalances epithelial growth to limit tissue overcrowding, Nature 484, 542 (2012).

[16] G. T. Eisenhoffer, P. D. Loftus, M. Yoshigi, H. Otsuna, C.-B. Chien, P. A. Morcos, and J. Rosenblatt, Crowding induces live cell extrusion to maintain homeostatic cell numbers in epithelia, Nature 484, 546 (2012).

[17] R. Levayer, C. Dupont, and E. Moreno, Tissue crowding induces caspase-dependent competition for space, Current Biology 26, 670 (2016).

[18] A. P. Le, J.-F. Rupprecht, R.-M. Mége, Y. Toyama, C. T. Lim, and B. Ladoux, Adhesion-mediated heterogeneous actin organization governs apoptotic cell extrusion, Nature Communications 12, 10.1038/s41467-020-20563-9 (2021).

[19] Y. Toyama, X. G. Peralta, A. R. Wells, D. P. Kiehart, and G. S. Edwards, Apoptotic force and tissue dynamics during drosophila embryogenesis, Science 321, 1683 (2008).

[20] G. M. Slattum and J. Rosenblatt, Tumour cell invasion: an emerging role for basal epithelial cell extrusion, Nature Reviews Cancer 14, 495 (2014).

[21] D. Boocock, N. Hino, N. Ruzickova, T. Hirashima, and E. Hannezo, Theory of mechanochemical patterning and optimal migration in cell monolayers, Nature Physics 17, 267 (2020).

[22] T. B. Saw, A. Doostmohammadi, V. Nier, L. Kocgozlu, S. Thampi, Y. Toyama, P. Marcq, C. T. Lim, J. M. Yeomans, and B. Ladoux, Topological defects in epithelia govern cell death and extrusion, Nature 544, 212 (2017).

[23] K. Kawaguchi, R. Kageyama, and M. Sano, Topological defects control collective dynamics in neural progenitor cell cultures, Nature 545, 327 (2017).

[24] G. Duclos, C. Blanch-Mercader, V. Yashunsky, G. Salbreux, J.-F. Joanny, J. Prost, and P. Silberzan, Spontaneous shear flow in confined cellular nematics, Nature Physics 14, 728 (2018).

[25] C. Blanch-Mercader, V. Yashunsky, S. Garcia, G. Duclos, L. Giomi, and P. Silberzan, Turbulent dynamics of epithelial cell cultures, Physical Review Letters 120, 10.1103/physrevlett.120.208101 (2018).

[26] T. H. Tan, J. Liu, P. W. Miller, M. Tekant, J. Dunkel, and N. Fakhri, Topological turbulence in the membrane of a living cell, Nature Physics 16, 657 (2020).

[27] J. Zhang, N. Yang, P. K. Kreeger, and J. Notbohm, Topological defects in the mesothelium suppress ovarian cancer cell clearance, APL Bioengineering 5, 036103 (2021).

[28] A. Doostmohammadi, S. P. Thampi, T. B. Saw, C. T. Lim, B. Ladoux, and J. M. Yeomans, Celebrating soft matter’s 10th anniversary: Cell division: a source of active stress in cellular monolayers, Soft Matter 11, 7328 (2015).

[29] A. Doostmohammadi, S. P. Thampi, and J. M. Yeomans, Defect-mediated morphologies in growing cell colonies, Physical Review Letters 117, 10.1103/physrevlett.117.048102 (2016).

[30] P. G. d. Gennes and J. Prost, The physics of liquid crystals, 2nd ed., The international series of monographs on physics No. 83 (Clarendon Press, Oxford, 1998) oCLC: 833446379.

[31] B. Loewe, M. Chiang, D. Marenduzzo, and M. C. Marchetti, Solid-liquid transition of deformable and overlapping active particles, Physical Review Letters 125, 10.1103/physrevlett.125.038003 (2020).

[32] W. T. M. Irvine, V. Vitelli, and P. M. Chaikin, Pleats in crystals on curved surfaces, Nature 468, 947 (2010).

[33] R. Mueller, J. M. Yeomans, and A. Doostmohammadi, Emergence of active nematic behavior in monolayers of isotropic cells, Physical Review Letters 122, 10.1103/physrevlett.122.048004 (2019).

[34] J. W. Cahn and J. E. Hilliard, Free energy of a nonuniform system. I. interfacial free energy, Journal of Chemical Physics 28, 258 (1958).

[35] B. Smeets, R. Alert, J. Pešek, I. Pagonabarraga, H. Ramon, and R. Vincent, Emergent structures and dynamics of cell colonies by contact inhibition of locomotion, Proceedings of the National Academy of Sciences 113, 14621 (2016).

[36] G. Peyret, R. Mueller, J. d’Alessandro, S. Begnaud, P. Marcq, R.-M. Mège, J. M. Yeomans, A. Doostmohammadi, and B. Ladoux, Sustained oscillations of epithelial cell sheets, Biophysical Journal 117, 464 (2019).

[37] M. C. Gibson, A. B. Patel, R. Nagpal, and N. Perrimon, The emergence of geometric order in proliferating metazoan epithelia, Nature 442, 1038 (2006).

[38] M. C. Marchetti, J. F. Joanny, S. Ramaswamy, T. B. Liverpool, J. Prost, M. Rao, and R. A. Simha, Hydrodynamics of soft active matter, Reviews of Modern Physics 85, 1143 (2013).

[39] A. Doostmohammadi, J. Ignés-Mullol, J. M. Yeomans, and F. Sagués, Active nematics, Nature Communications 9, 10.1038/s41467-018-05666-8 (2018).

[40] D. Wenzel and A. Voigt, Multiphase field models for collective cell migration, arXiv preprint arXiv:2106.10552 (2021).

[41] M. Bär, R. Großmann, S. Heidenreich, and F. Peruani, Self-propelled rods: Insights and perspectives for active matter, Annual Review of Condensed Matter Physics 11, 441 (2020).

[42] O. J. Meacock, A. Doostmohammadi, K. R. Foster, J. M. Yeomans, and W. M. Durham, Bacteria solve the problem of crowding by moving slowly, Nature Physics 17, 205 (2020).

[43] A. Amiri, R. Mueller, and A. Doostmohammadi, Half-integer and full-integer topological defects in polar active matter: Emergence, crossover, and coexistence, arXiv:2106.03144 [cond-mat, physics:physics] (2021), arXiv: 2106.03144.

[44] J. Christoffersen, M. Mehrabadi, and S. Nemat-Nasser, A micro-mechanical description of granular material behavior, Journal of Applied Mechanics 48, 339 (1984).

[45] H. Li, D. Matsunaga, T. S. Matsui, H. Aosaki, K. Inoue, A. Doostmohammadi, and S. Deguchi, Wrinkle force microscopy: a new machine learning based approach to predict cell mechanics from images (2021), arXiv:2102.12069 [physics.bio-ph].

[46] T. S. Majmudar and R. P. Behringer, Contact force measurements and stress-induced anisotropy in granular materials, Nature 435, 1079 (2005).

[47] D. T. Tambe, C. C. Hardin, T. E. Angelini, K. Rajendran, C. Y. Park, X. Serra-Picamal, E. H. Zhou, M. H. Zaman, J. P. Butler, D. A. Weitz, J. J. Fredberg, and X. Trepat, Collective cell guidance by cooperative intercellular forces, Nature Materials 10, 469 (2011).

